# The Life Span of the Great Black Wasp *Sphex pensylvanicus* Linnaeus, 1763 (Hymenoptera: Sphecidae) and Other Observations from a Nesting Aggregation in Sioux City, Iowa, USA. 2003-2017

**DOI:** 10.1101/2022.01.14.476354

**Authors:** G.K. Lechner

## Abstract

In a fifteen year period, the observed life span of both male and female *Sphex pensylvanicus* was documented in an aggregation of these wasps behind a retaining wall on a residential lot in Sioux City, Iowa, USA. In addition, early season emergence of the males before the females (protandry) and instances of male territorial behavior were also confirmed. Half or more of the wasps disappeared; possible out-migration?

## Introduction

The Great Black Wasp, *Sphex pensylvanicus* L., is a solitary fossorial sphecid wasp that ranges over much of the continental United States (Bohart and Menke, 1963) and portions of southern Canada (Buck, 2004). According to an even more recent report, *S. pensylvanicus* has now been found in southern New Brunswick, Canada (Lewis, 2020). It is the largest sphecid wasp in eastern North America except for the Cicada Killer, *Sphecius speciosus* (Kurczewski, 1997). Although classified as solitary, female *Sphex pensylvanicus* are known to nest in aggregations, seemingly taking no interest in each other, and preferring to nest in sheltered places such as tool sheds, green houses and storm drains (Frisch, 1938; Rau, 1944; Kurczewski, 1997). Previous authors have also reported female *S. pensylvanicus* in the same aggregation sites for at least three consecutive years (Rigley and Hays, 1977).

In this paper, I set forth some of my observations of *Sphex pensylvanicus* from an aggregation that had been in existence for at least fifteen years on a residential lot in Sioux City, Iowa, and data has been collected regarding this aggregation from 2003 through 2017. I have previously published shorter papers on wasps from this aggregation regarding homing flight distance challenges (Lechner, 2015) and prey interactions (Lechner, 2016).

## Observations

The observation site is my residential lot (with single family dwelling and detached one car garage) in the 3600 block of Virginia St., Sioux City, Iowa. This block is on a hill; and to accomodate the rise of the hill, lots on the upper reaches of the block are terraced leaving each lot relatively level.

Terracing necessitated the construction of retaining walls along each lot line. My lot at 3624 Virginia St. Is approximately five feet (approx. 1.5 m) higher than the lot to the south and approximately five feet lower than the lot to the north. The North Lot Line Retaining Wall (NLLRW) is 177 feet long (approx. 53.9 m) and is built of standard 8” x 8” x16” (approx. 20 × 20 × 40 cm) concrete blocks for approximately ¾ of its length beginning at the street (Fig. 1) and railroad ties for the remainder of its length to the rear of the lot (Fig. 2) and is capped with poured concrete. This wall supports the poured concrete driveway at 3628 Virginia along its entire length.

**Figure 1.**
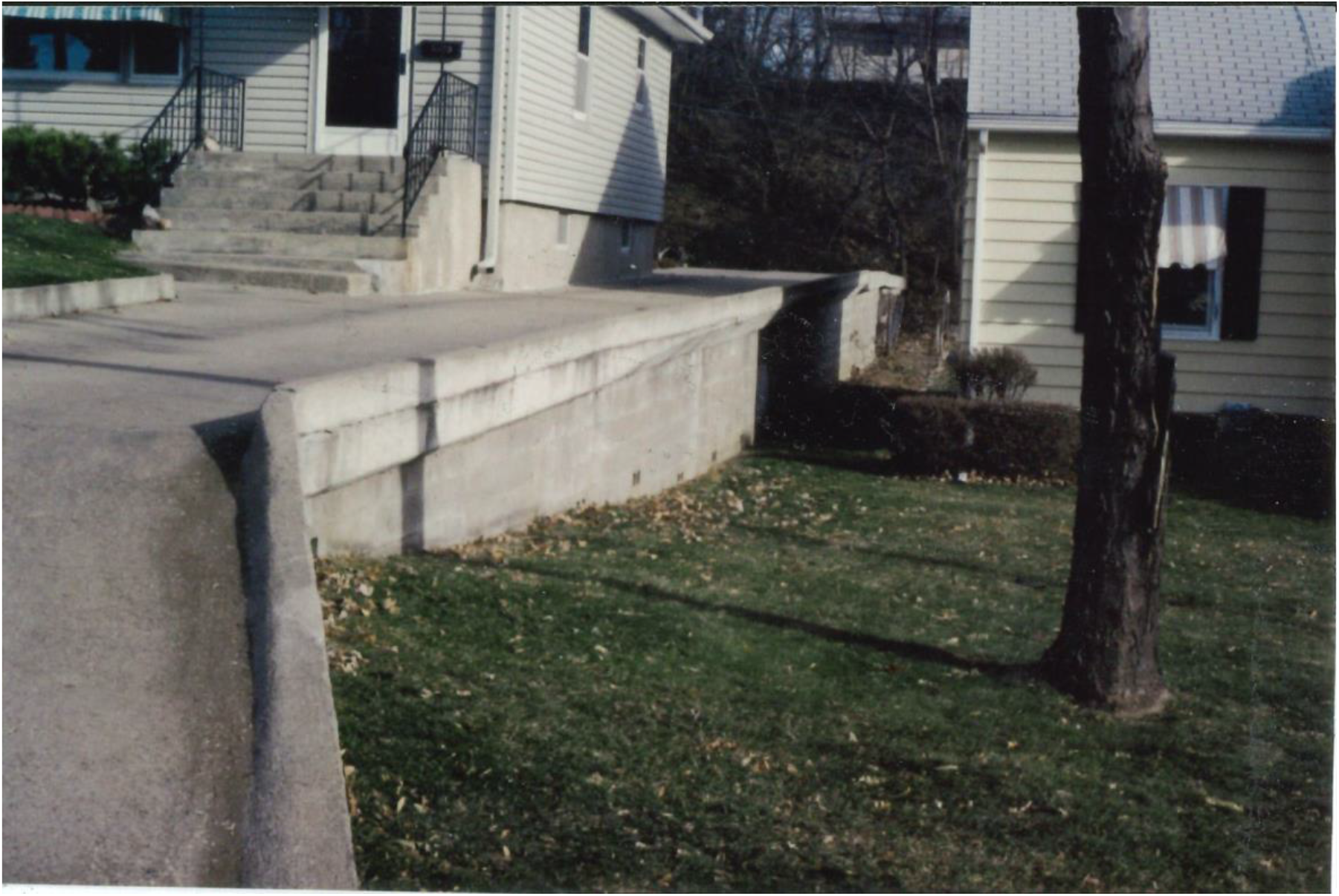
A view of the North Lot Line Retaining Wall at 3624 Virginia St., Sioux City, Iowa, looking east This photo taken in 2003.

**Figure 2.**
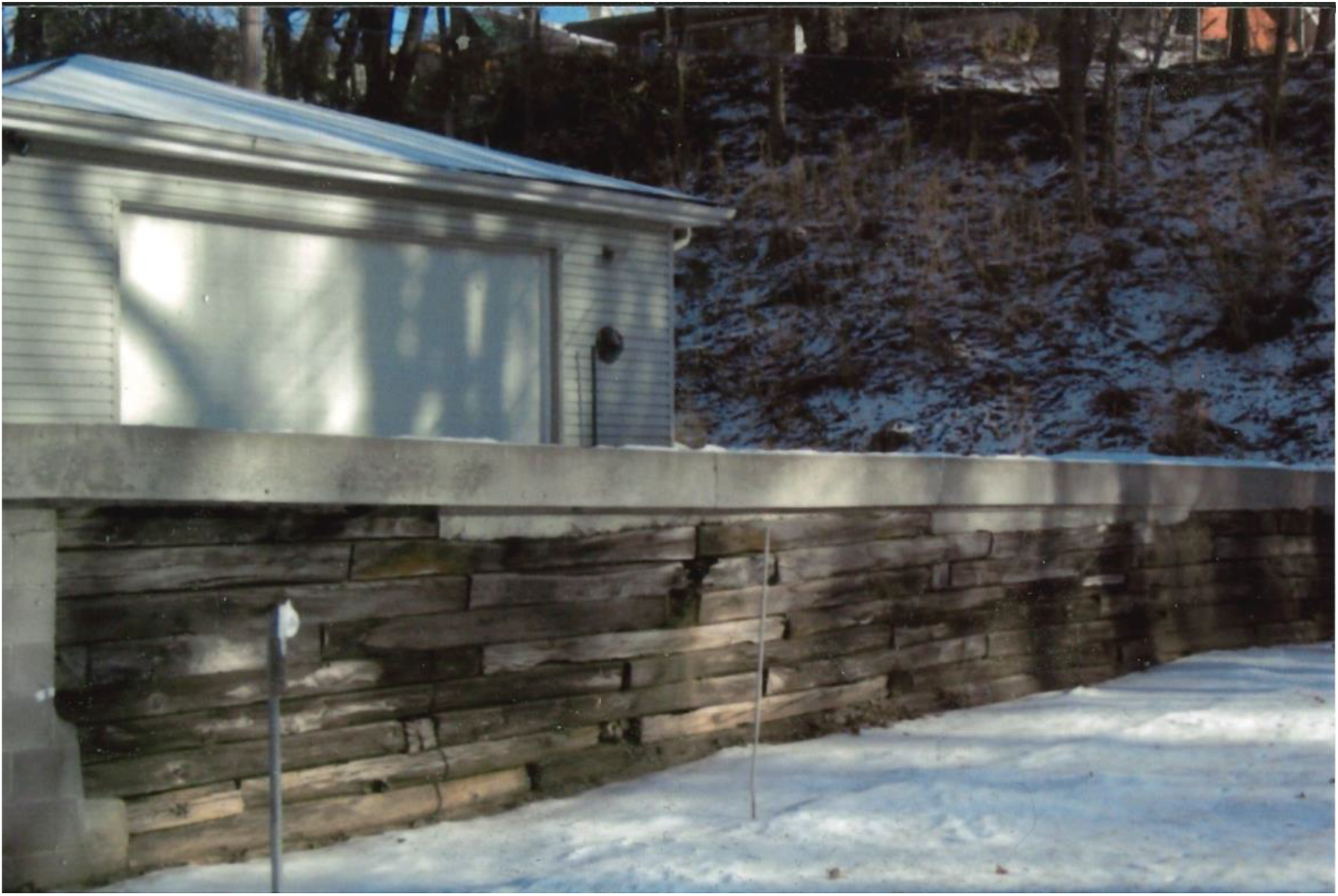
The railroad tie portion of the North Lot Retaining Wall, 3624 Virginia St., Sioux City, Iowa, This photo taken in winter 2013.

When the NLLRW was built, ten of the concrete blocks in the lower courses of the wall were installed sideways (i.e. rotated 90° on the long axis) for water drainage/pressure relief thereby leaving the usually concealed hollows inside the blocks open to the environment. These exposed hollows have served as apertures that have been attractive to fossorial wasps and function as common entryways (i.e. as a sort of atrium or vestibule) to their burrows in the soil behind the wall. For purposes of description, I have numbered these ten blocks 1 through 10 from west to east, and designated the apertures either E (for east) or W (for west). This numbering system should become clear by referring to Figure 1 and to the Elevation Diagram (Figure 3). I will use this block and aperture numbering system throughout this report. Infrequently, Cicada Killers (*Sphecius speciosus*), Steel Blue Cricket Hunters (*Chlorion aerarium*), and even more rarely, Black and Yellow Mud Daubers (*Sceliphron caementarium*) and *Liris argentatus* have visited the apertures of the NLLRW. However, *Sphex pensylvanicus* was by far the most common nester in the NLLRW although the dates of first appearence have been variable – as early as 20 June (in 2004) and as late as 20 July (in 2010). Consequently, the season length of *S. pensylvanicus* activity in the NLLRW was also variable – as long as 84 days in 2004 and as short as 21 days in both 2010 and 2014 (See Table 1).

**Table 1.** Dates of Observations of *Sphex pensylvanicus* L. Anywhere on the Lot and at the North Lot Line Retaining Wall, 3624 Virginia St., Sioux City, Iowa, 2003 – 2017.

| Year | Anywhere on Lot |  |  | North Lot Line Retaining Wall |  |  |
| --- | --- | --- | --- | --- | --- | --- |
|  | From | Through | Season Length (Days) | From | Through | Season Length (Days) |
| 2003 | 4 July | 17 Sept | 76 | 4 July | 17 Sept | 76 |
| 2004 | 20 June | 11 Sept | 84 | 20 June | 11 Sept | 84 |
| 2005 | 7 July | 23 Aug | 48 | 7 July | 23 Aug | 48 |
| 2006 | 3 July | 26 Aug | 55 | 3 July | 26 Aug | 55 |
| 2007 | 30 June | 9 Aug | 41 | 3 July | 7 Aug | 36 |
| 2008 | 1 July | 1 Sept | 63 | 1 July | 24 Aug | 55 |
| 2009 | 3 Aug | 19 Sept | 49 | 14 Aug | 19 Sept | 37 |
| 2010 | 8 July | 9 Aug | 33 | 20 July | 9 Aug | 21 |
| 2011 | 11 July | 28 Aug | 49 | 11 July | 28 Aug | 49 |
| 2012 | 24 June | 13 Aug | 51 | 24 June | 13 Aug | 51 |
| 2013 | 24 July | 11 Sept | 50 | 24 July | 11 Sept | 50 |
| 2014 | 1 July | 2 Sept | 64 | 2 Aug | 22 Aug | 21 |
| 2015 | 18 July | 16 Aug | 30 | 18 July | 16 Aug | 30 |
| 2016 | 1 July | 15 Aug | 37 | 24 July | 15 Aug | 23 |
| 2017 | 7 July | 30 Aug | 55 | 8 July | 30 Aug | 54 |

**Figure 3.**
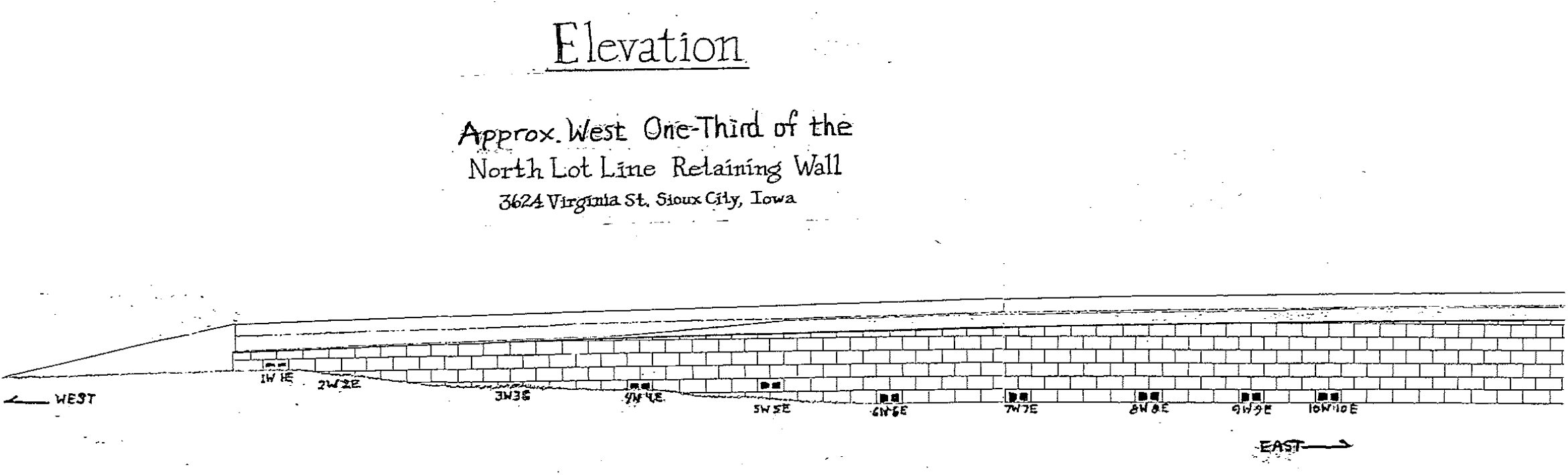
Elevation Diagram of the Approximate West One-Third of the North Lot Line Retaining Wall, 3624 Virginia St., Sioux City, Iowa.

### Observational Methods

For most of the study I monitored the activities of *Sphex pensylvanicus* at the study site in two ways. One method involved random walks around the lot sporadically. The other method was by sitting at an observation post, quite frequently at the NW corner of the house, to keep the NLLRW in view.

A sweep net was used to snare the flying *S. pensylvanicus* or a trap jar would be set in the apertures to catch the wasps on their emergence. If a captured wasp was to be marked, it was brought indoors and placed in a freezer for usually no more than five minutes until “anesthetized”. While in this stuperous state, a mark of paint would be placed on the scutum. Alternatively, instead of paint, a small numbered tag would be glued on the scutum with cyanoacrylate glue.

In 2003 when this project began, I intended to mark each captured wasp with a unique paint color and design. It quickly became apparent that there were going to be too many wasps for this painting system to be workable. Therefore, I switched to the numbered tag system. Since *Sphex pensylvanicus* does not overwinter in the adult stage, I could start over each season with tag #1 and identify each wasp consecutively by order of capture.

### Observational Problems

It seems appropriate at this point to discuss some of the problems that I encountered in this project. First, much of the literature on fossorial wasps includes information, photos and diagrams about excavation of nest sites (e.g. Evans, 1966b; O’Neill, 2001). From the photos of the NLLRW (Figures 1 and 2), one can see that excavation at this study site is impossible; the NLLRW is owned by the neighbors at 3628 and holds up their driveway. So, regrettably my findings lack such data as length or depth of the wasps’ tunnels to their nests, condition of the larvae or cocoons, etc.

In 2003, the first year of the study, I found that the round trap jars inserted into the block apertures did not fit tightly enough to preclude the wasps’ evading capture by bypassing the jars. To eliminate this problem, before the 2004 season, I fitted each aperture (except in Blocks 2 and 3) with a plywood collar so that the full area of the aperture was covered. A round hole matching the diameter of a trap jar was cut through the center of the collar into which the jar could be fit tightly. In this way no wasp could evade the trap when exiting the aperture nor could a returning wasp enter unless I removed the jar first. Interestingly, over the many years of this study, the number of times returning females found their apertures blocked by a trap jar were in the multiple dozens; and rather than fly away, frequently they would hover near the block until I approached and removed the trap jar, then fly into the aperture. It was almost as if they were waiting for me to remove the impediment so they could enter. For clarification refer to Figure 4, a photograph of Block 5 showing a trap jar fitted into the 5E aperture whereas the 5W aperture is open.

**Figure 4.**
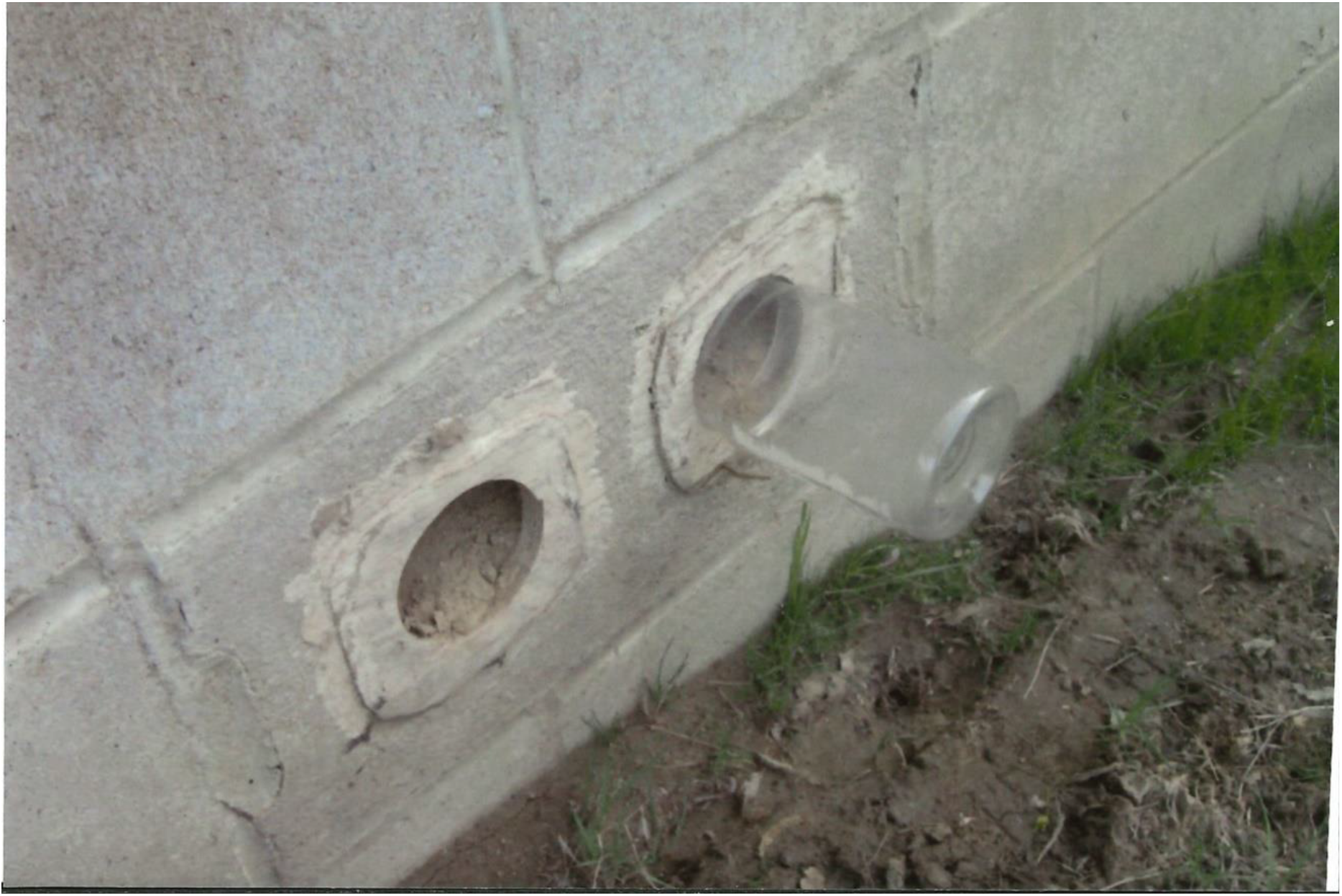
A photo of Block 5 in the North Lot Retaining Wall. Note the trap jar inserted into the 5E aperture (right) whereas the 5W aperture (left) is open to the environment.

To continue the discussion of difficulties, consider that to observe all activities of a diurnal wasp like *Sphex pensylvanicus* would require an observer to be present at the nesting site from dawn to dusk. This was not possible. Although in some years of the study continuous observation of the NLLRW amounted to many hours (e.g. over 137 hours in 2003; over 112 hours in 2004) yielding over 2000 separate sightings of *S. pensylvanicus*, the wall was not watched at all times; and on some days the study site was not visited at all. Additionally, every time I took an unmarked wasp indoors to be given identification, a further absence of at least fifteen minutes would have occurred.

I should also mention that my observational time was not consistent from season to season. As stated above, in some years I was able to devote many hours to the project. In other years, other obligations limited my time. For instance, in 2009 I made about thirty sightings of *S. pensylvanicus* yet only one wasp was marked that year (See Lechner, 2016). Also, in another example, in 2011, sightings of *S. pensylvanicus* exceeded one hundred, but I only marked five wasps that year.

Furthermore, even if an observer were available to watch the NLLRW from sunrise to sunset, the length of the wall encompassing the distance from Block 1 to Block 10 (approx. 70 feet) is too large for one person to see every entry and exit of a wasp from these apertures. For example, if one’s attention is turned to wasp activity at Block 1, anything happening near Block 10 could be missed easily. In other words, the wall is just too long for one pair of eyes to see everything. Still, that being said, from 2003 through 2017, as noted above, many sightings were made. Obviously with a nesting aggregation, sightings could have been of the same wasp multiple times. For example, in 2004, the tagged wasp #63 was recaptured 32 times between 21 July and 12 August. Another example is Wasp #16 from 2012. She was encountered 41 times between 13 July and 13 August of that year.

Another hinderance to easy observation is the wasps themselves. The females are fast and evasive fliers; and, unless feeding on flowers, are rarely found sitting still long enough to read their tag number or see a paint mark. So identification usually required a recapture, sometimes by net, but most often by trap jar in the apertures of the NLLRW.

I should also mention here that I did not turn my attention to the railroad tie portion of the NLLRW (Fig. 2) until 2014 when it seemed that the *Sphex pensylvanicus* had ceased their activities in the concrete block portion of the NLLRW, and in that case I moved my observation spot to the back yard so as to be closer to the railroad ties.

Lastly, the final insult to this aggregation of *S. pensylvanicus* occurred in early June of 2018 when the owners of the NLLRW had the entire length of the wall plastered over blocking some of the apertures (Fig. 5) Only Blocks 5, 7, 8, 9 and 10 still have exposed apertures. I have not seen any wasp activiity at the NLLRW since 2017.

**Figure 5.**
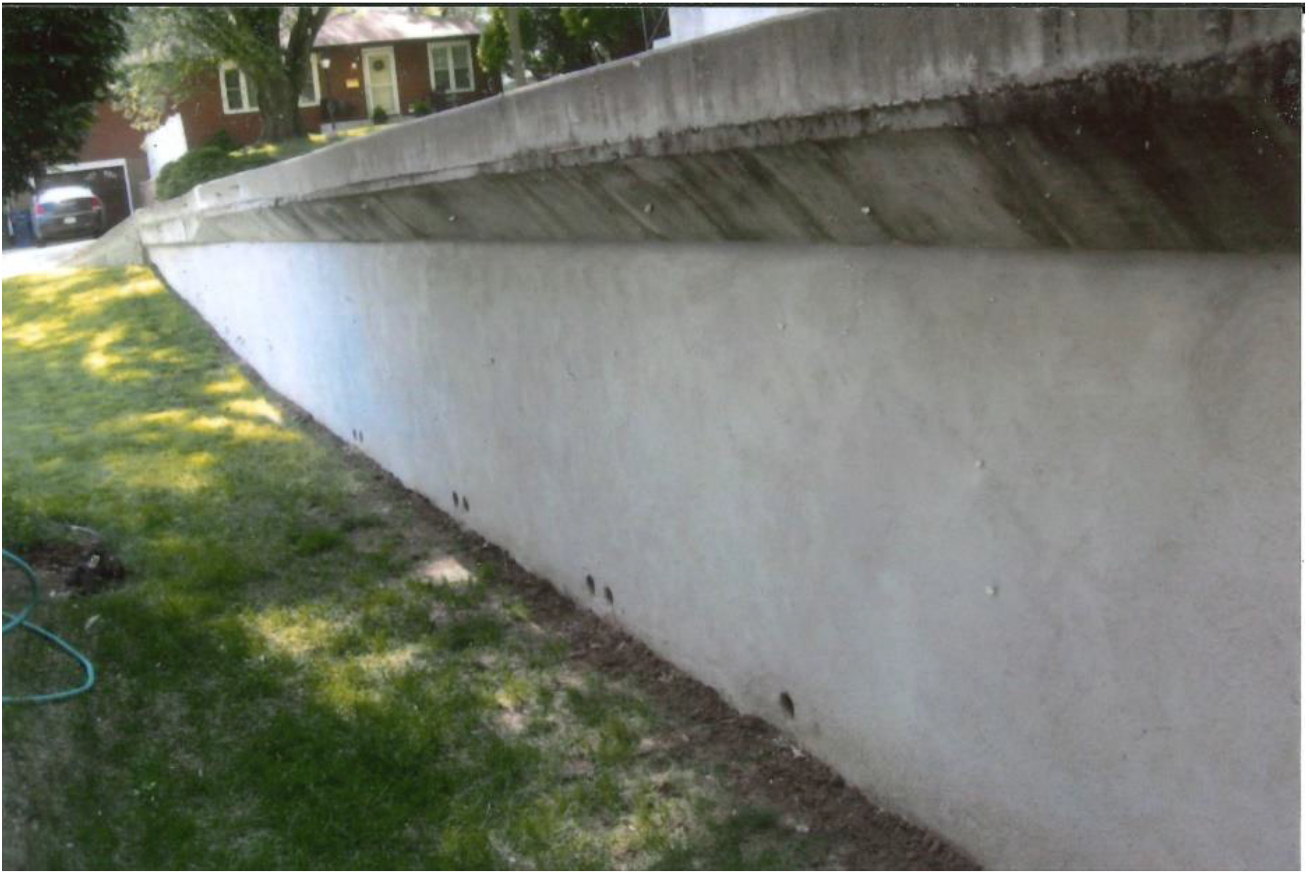
A photo of a portion of the North Lot Line Retaining Wall looking west after the wall had been plastered over in June 2018.

## Observed Life Span of *Sphex pensylvanicus*

Rau (1944) observed activities of *S. pensylvanicus* in an aggregation of these wasps at a location near St. Louis, Missouri, in 1938 and 1939; he reported an adult life span of over three months. In all likelihood, this is an over estimation. Rau may have overlooked the probability of two generations per season.

More recently, in a comprehensive paper regarding a nesting aggregation of *Sphex pensylvanicus* in New York State, F.E. Kurczewski (1997) reported that only three of 22 wasps lived longer than a month with a maximum of 35 days.

At my study location, from 2003 through 2017, I marked 417 adult *S. pensylvanicus*. Many of these wasps were only seen one day (i.e. the day of their initial capture and marking). Nevertheless, throughout the entire length of this study, I noted that a number of wasps were alive a remarkably long time. The female wasp with the longest confirmed life span (37 days) was Wasp #74 from 2004. She was first captured on 21 July and last captured on 1 September. Wasp #74 was also recaptured thirteen times in that 37 day span. If female longevity were linked to robust body size, Wasp #74 would belie the correlation. At 25 mm in length, she was among the more petite females encountered during the study (See Table 2).

**Table 2.** Body Length Measurements of 331 *Sphex pensylvanicus* L. From the Study Site, 3624 Virginia St., Sioux City, Iowa. 2003 – 2017.

|  | Number of Wasps | Average (mm) | Range (mm) |
| --- | --- | --- | --- |
| ♂ | 138 | 21.2 | 18-27 |
| ♀ | 193 | 29.0 | 19-34 |

The next longest observed life span (33 days) was accorded to Wasp #5 from the 2003 season. She was first captured on 15 July and last encountered on 16 August yet she was only seen four times between those two dates.

The third longest observed life span (32 days) goes to Wasp #16 from 2012. She was first caught on 13 July and last on 13 August. This wasp was a very busy nester as she was encountered 41 times between these two dates.

Regarding the life span of males, Kurczewski (1998) reported on nine males he observed at a nesting aggregation of *S. pensylvanicus* females in a storm drain at Marcellus, New York, in 1982. He reported individual male life spans of only 10-14 days with just three of the males living an entire two week period. As to the male *S. pensylvanicus* found at my study site, I can confirm some male longevity that exceeds this handsomely.

I found two males whose confirmed life span was at least twenty days. Wasp #14 from 2003 was captured and tagged on 20 July and was last seen on 12 August. Between those two dates, he was encountered 43 more times, and there were nine days in that stretch of time where I did not see him at all. He had established a territorial post on a sprig of ground level plant on the <u>south</u> lot line retaining wall, and I encountered him there frequently as I walked by this location (See Territoriality).

Another long lived male was Wasp #100 from 2004. He was captured on 14 August as he was feeding on flowers of mint (Menthus spp.) growing in the basement window well on the east side of the house. I walked by this location multiple times daily, and Wasp #100 seemed to be there almost constantly. He was seen on 107 separate occasions for the 27 days, except for five, through 9 September, each time feeding on the mint flowers.

For a summary of female and male *Sphex pensylvanicus* longevity throughout the length of this study, refer to Tables 3 and 4.

**Table 3.** Observed Life Span (Days) of Marked Female *Sphex pensylvanicus*, 3624 Virginia St. Sioux City, Iowa. 2003-2020

| Year | No. of Wasps seen Only One Day | No. of Wasps seen Multiple Days | Wasps with the Two Longest Observed Life Spans |  |  |  |  |
| --- | --- | --- | --- | --- | --- | --- | --- |
|  |  |  | Identifi-<br>cation<br>(# or<br>paint<br>color) | Observed<br>Life Span<br>(Days) | From | Through | Number of<br>Encounters |
| 2003 | 29 | 33 | #5 | 33 | 15 July | 16 Aug | 5 |
|  |  |  | "1-Blue Dot" | 30 | 9 July | 7 Aug | 3 |
| 2004 | 27 | 37 | #74<sup>1</sup> | 37 | 21 July | 1 Sept | 14 |
|  |  |  | #58 | 31 | 18 July | 17 Aug | 13 |
| 2005 | 3 | 1 | #2 | 3 | 13 July | 15 July | 2 |
|  |  |  | #7 | 1 | 2 Aug | - | 1 |
| 2006 | 18 | 5 | #41 | 19 | 18 July | 5 Aug | 5 |
|  |  |  | #49 | 17 | 27 July | 12 Aug | 8 |
| 2007 | 3 | 9 | #8 | 16 | 19 July | 3 Aug | 5 |
|  |  |  | #6 | 13 | 12 July | 24 July | 4 |
| 2008 | 11 | 6 | #29 | 24 | 1 Aug | 24 Aug | 11 |
|  |  |  | #30 | 13 | 1 Aug | 13 Aug | 9 |
|  |  |  | #35 | 13 | 12 Aug | 24 Aug | 5 |
| 2009 | - | 1 | "White"<sup>2</sup> | 8 | 1 Sept | 8 Sept | 6 |
| 2010 | 1 | 1 | "White" | 7 | 1 Aug | 7 Aug | 7 |
|  |  |  | "Yellow" | 1 | 9 Aug | - | 1 |
| 2011 | 1 | 4 | "White" | 18 | 31 July | 17 Aug | 15 |
|  |  |  | "Yellow" | 7 | 17 July | 23 July | 13 |
| 2012 | 15 | 13 | #16 | 32 | 13 July | 13 Aug | 41 |
|  |  |  | #11<sup>3</sup> | 24 | 7 July | 30 July | 8 |
| 2013 | 5 | 2 | #14<sup>4</sup> | 3 | 31 July | 1 Aug | 3 |
|  |  |  | #19<sup>5</sup> | 2 | 20 Aug | 21 Aug | 2 |
| 2014<sup>6</sup> | 2 | 6 | #3 | 20 | 2 Aug | 21 Aug | 3 |
|  |  |  | #8 | 6 | 17 Aug | 22 Aug | 3 |
| 2015<sup>7</sup> | 3 | 8 | #2 | 20 | 19 July | 7 Aug | 11 |
|  |  |  | #1 | 9 | 18 July | 26 July | 8 |
| 2016 | 3 | 5 | #6 | 18 | 27 July | 13 Aug | 8 |
|  |  |  | #3 | 8 | 24 July | 31 July | 6 |
| 2017 | 7 | 1 | "Brown" | 7 | 8 Aug | 14 Aug | 3 |
| 2018 | 0 | 0 | No *Sphex pensylvanicus* observed this season |  |  |  |  |
| 2019 | 0 | 0 | No *Sphex pensylvanicus* observed this season |  |  |  |  |
| 2020 | 0 | 0 | No *Sphex pensylvanicus* observed this season |  |  |  |  |
| Totals | 128 | 132 |  |  |  |  |  |

1. This wasp had the longest observed life span throughout the length of the study. 2. This wasp was the only one marked this season even though about a dozen sightings of *S. pensylvanicus* were made. For interactions with prey, (see Lechner, 2016). 3. This wasp was visiting the flowers of mint in the basement window well on 28 and 30 July. 4. This wasp was given a homing flight return challenge on 1 August from which she was not seen again. 5. This wasp was from the railroad tie portion of the NLLRW. 6. and 7. All wasps from these two years were from the railroad tie portion of the NLLRW.

**Table 4.** Observed Life Span (Days) of Marked Male *Sphex pensylvanicus*. 3624 Virginia St. Sioux City, Iowa. 2003 – 2020.

| Year | No. of Wasps seen Only One Day | No. of Wasps seen Multiple Days | Wasps with the Two Longest Observed Life Spans |  |  |  |  |
| --- | --- | --- | --- | --- | --- | --- | --- |
|  |  |  | Identification (# or paint color) | Observed Life Span (Days) | From | Through | Number of Encounters |
| 2003 | 8 | 11 | #14 | 24 | 20 July | 12 Aug | 44 |
|  |  |  | "1-Green Line" | 19 | 12 July | 22 July | 20 |
| 2004 | 38 | 8 | #100 | 27 | 14 Aug | 9 Sept | 107<sup>1</sup> |
|  |  |  | #1<sup>2</sup> | 6 | 20 Jun | 25 June† |  |
| 2005 | 1 | 0 | #4 | 1 | 31 July | - | 1 |
| 2006 | 32 | 2 | #22<sup>3</sup> | 3 | 10 July | 12 July† |  |
|  |  |  | #11<sup>4</sup> | 2 | 7 July | 8 July† |  |
| 2007 | 2 | 1 | #3<sup>5</sup> | 2 | 7 July | 8 July |  |
| 2008 | 20 | 1 | #37<sup>6</sup> | 14 | 17 Aug | 30 Aug | 7 |
| 2009 | - | - |  |  |  |  |  |
| 2010 | - | - |  |  |  |  |  |
| 2011 | - | - |  |  |  |  |  |
| 2012 | 1 | 0 | #18<sup>7</sup> | 1 | 14 July | - | 1 |
| 2013 | 12<sup>8</sup> | 0 |  |  |  |  |  |
| 2014 | - | - |  |  |  |  |  |
| 2015 | - | - |  |  |  |  |  |
| 2016 | - | - |  |  |  |  |  |
| 2017 | 20<sup>9</sup> | - |  |  |  |  |  |
| 2018 | 0 | 0 | No *Sphex pensylvanicus* observed this season |  |  |  |  |
| 2019 | 0 | 0 | No *Sphex pensylvanicus* observed this season |  |  |  |  |
| 2020 | 0 | 0 | No *Sphex pensylvanicus* observed this season |  |  |  |  |
| Totals | 134 | 23 |  |  |  |  |  |

1. All sightings made while wasp was visiting window well mint flowers. 2. This wasp emerged unhealthy. Only seen for six days because it was kept indoors and fed sugar water in an attempt at resuscitation. 3. Unhealthy and deformed. Kept indoors and fed sugar water. 4. Kept in a jar overnight. Dead the next day. 5. Lethargic. Brought indoors and fed sugar water. Released the next day by placing on mint plants. It did not fly away but was gone the next day. 6. All sightings of this wasp were on the flowers of mint at the NE corner of the lot. 7. A confirmed emerger from the NLLRW – Aperture 6W. 8. All males in 2013 seen only one day. 9. All males in 2017 seen only one day. One died, probably from overheating in a trap jar.

### Protandry

Evans (1966b) reported that in nearly all solitary wasps and bees, the males emerge several days before the females. Beginning with the 2004 season, I was able to confirm this phenomenon by installing trap jars in most of the apertures of the NLLRW before the season’s first emergence of *Sphex pensylvanicus*,

The installation of pre-emergence trap jars occurred in 2004, 2006, 2007, 2008, 2013, 2014, 2015, 2017, 2018, 2019, 2020 (See Table 5). The table shows that at my study site, males generally appeared earlier than females as predicted by previous authors. First emergence occurred as early as 20 June (in 2004) and as late as 24 July (in 2013). For example, in 2004 from 20 June through 11 July 43 males were emergent from the apertures of the NLLRW whereas only two females were caught in this time frame. Then, later in the year, from 13 July through 5 September, there were no males trapped from the apertures yet 41 females were captured in this latter part of the season.

**Table 5.** Numbers of Observed Male and Female *Sphex pensylvanicus* in the First Twenty Days of the Season Compared with Numbers of Observed Male and Female *Sphex pensylvanicus* in the Remainder of the Season. 2003 -2020.

| Year | Pre-emergence Traps Jars Installed in Apertures of the NLLRW (Date) | No. of Wasps Seen in First 20 Days of Season |  | No. of Wasps Seen in Remainder of Season |  | Comments |
| --- | --- | --- | --- | --- | --- | --- |
|  |  | ♂ | ♀ | ♂ | ♀ |  |
| 2003 | - | ~ 20 | 21 | 8 | 38 |  |
| 2004 | 14 June | 38 | 1 | 8 | 63 |  |
| 2005 | - | 0 | 2 | 1 | 2 | No preemergence traps set this year. About 60 confirmed sightings of *S. pen.* but only five wasps marked |
| 2006 | 7 June | 33 | 12 | 1 | 11 |  |
| 2007 | 14 June | 3 | 5 | 0 | 7 |  |
| 2008 | 16 June | 6 | 2 | 15 | 15 |  |
| 2009 | - |  |  |  |  | No preemergence traps set this year. Approx. 30 *S.pen.*sightings but only one wasp marked. |
| 2010 | - |  |  |  |  | No preemergence traps set this year. Approx. 20 *S.pen.* sightings. Season length was only 21 days |
| 2011 | - |  |  |  |  | No preemergence traps set this year. Well over 100 *S. pen.* sightings. |
| 2012 | - | 0 | 15 | 1 | 24 | No preemergence traps set this year. |
| 2013 | 13 June | 12 | 6 | 0 | 1 | The one female from the latter part of the season emerged from the railroad tie portion of the NLLRW. |
| 2014 | 14 June | - | - | - | 8 | Preemergence traps were set, but no wasps emerged from the apertures. The 8 females emerged from the railroad ties. |
| 2015 | 17 June | - | - | - | 11 | No emergers from the concrete block apertures. All 11 females from the railroad tie portion of the NLLRW. |
| 2016 | - | 0 | 7 | 0 | 1 | No preemergence traps set this year. No males seen. All females emerged from the concrete block apertures. |
| 2017 | 7 June | 20 | 3 | 0 | 4 | All wasps (except for 4 from the railroad ties) emerged from the concrete block apertures. |
| 2018 | 15 June | - | - | - | - | No wasp activity. The entire NLLRW was plastered over leaving only Blocks 5,7,8,9 and 10 with exposed apertures. |
| 2019 | 11 June | - | - | - | - | No wasp activity this year |
| 2020 | 11 June | - | - | - | - | No wasp activity this year. |

Likewise, in 2017, the first year in which *S. pensylvanicus* males emerged from the concrete blocks after a three year absence, 20 males emerged between 8 July and 24 July with only one female caught in that time span; and from 25 July through 8 August no males emerged.

### Territoriality

The literature has many reports of the territoriality of male solitary wasps (e.g. Davis, 1920; Lin, 1963; Evans, 1966 a,b; and especially O’Neill, 2001, for a comprehensive review). Such behavior has also been observed and discussed for male *Sphex pensylvanicus* (Kurczewski, 1998).

I can also add my observations of male *S. pensylvanicus* behaving in a territorial manner, and it was in 2003 that this behavior was most abundant. On the NLLRW, the males were noted to frequent one location and challenge any wasp approaching them. They would rest on the wall above the block apertures, sometimes flying down and landing on the lawn just in front of these entrances; and on occasions making seemingly unprovoked departures out of sight for some time only to return later and resume their watchful behavior.

The males were by no means quiet while on this sentinal-like duty but behaved like skittish busybodies. Without any obvious cause, the male would fly upward from his station and then land just seconds later. A bird flying by, even a passing car in the street, could set him in motion.

The arrival of another wasp elicited an immediate confrontation. Both wasps would begin aerial “dogfighting”, flying quickly upward in a very tight intertwining spiral pattern. On two occasions in 2003 (16 and 17 July), I saw a trio of wasps performing in this manner. After flying upward, they would dive back down nearly to ground level, but not land, then repeat their upward spiraling. This went on for over two minutes in one instance and almost five minutes in the other. Then again on 20 and 26 July 2003, five more instances of three wasps making these spiraling flights were observed.

As examples of individually marked males exhibiting territorial behavior, I first offer Wasp #3 from 2003. Initially captured on 14 July, he was recaptured three times and was seen an additional eleven times (including four separate occasions on 26 July) while either resting, flying, or “dogfighting” other wasps in the proximity of Block 1.

Wasp #3 did show some tolerance at times for the presence of another male. On both 16 and 18 July 2003, Wasp #3 was noted on the wall above Block 1 with the wasp designated as “Green Line” (for the paint design I put on his scutum). “Green Line” was present here from initial capture on 12 July 2003 to his last sighting on 22 July 2003.

“Green Line” was noted to pay exceedingly close attention to the 1W aperture in Block 1. On 20, 21 and 22 July 2003, he was observed to have stationed himself on the vertical surface of the NLLRW only two inches or less above the aperture; and in three instances, I found that he had caught himself in the trap jar. As mentioned previously, in 2003 the trap jars did not fit snugly in the apertures so this wasp could have easily found his way into the aperture around the trap jar then have caught himself trying to make an exit (See Observational Problems).

Another territorial male was Wasp #13 first captured on 20 July 2003. He was recaptured four times and sighted another four times between that date and 28 July. He had set up his territory in the area of the NLLRW a few feet above Blocks 5 and 6. The same resting and flying patterns were exhibited by Wasp #13.

Also in 2003, Wasp #14 was a male first captured on 20 July from his territorial station on the top of the <u>south</u> lot line retaining wall. While I did not engage in any lengthy continuous observation of this location, I walked by many times daily, and Wasp #14 was seen resting on vegetation, “dogfighting” other wasps (including a Steel Blue Cricket Hunter, *Chlorion aerarium*, on 8 August), or flying lazily around the area. Between 20 July and 12 August, when he was last encountered, I recaptured Wasp #14 nine times and sighted him an additional 34 times.

In the next season, 2004, only one instance of possible male territorial behavior was encountered. On 10 July of that year, a male was found resting on the vertical surface of the NLLRW above Block 10. He was captured, and I affixed tag #56 to his scutum. Upon release, he did not fly away but rested in the lawn below Block 10. About 45 minutes later, Wasp #56 was still there. I picked him up and tossed him into the air, and he flew upward and north. Wasp #56 was never seen again.

In 2005, again only one instance of possible male territorial behavior presented itself. On 31 July of that year, I found a male sitting on the vertical surface of the NLLRW approximately 18 inches above Block 4. This wasp was given tag #4, and after the tagging procedure, was released and never reencountered.

Interestingly, from 2006 through 2017, and even though the females were nesting in the NLLRW during most of that time frame, no further instances of possible male territorial behavior were seen anywhere on the NLLRW nor anywhere else for that matter.

## Discussion

During the years that I have been observing the activities of *Sphex pensylvanicus* on my residential lot, a number of situations have arisen which present questions. I will discuss these in the following subsections.

### Longevity

From Tables 3 and 4, note that of all the wasps I marked for identification, many were never reencountered after their initial capture. Breaking this down by gender, 53 of 76 males (∼70%) and 128 of 260 females (∼49%) were lost. Even taking into account my non-continuous observational methods (See Observational Problems), this seems like a very high number of wasps to apparently just disappear. Did they fly away and die or did they emigrate from this study site seeking nesting opportunities elsewhere? Polidori et.al. (2010) reported on marked males (n=27) and females (n=90) of the digger wasp *Stizus continuus* from a study done over three different years at a location in Spain. A somewhat similar percentage of “Lost” was found in that study – 78% for males and 50% for females. Dr. Polidori has informed me that mortality alone probably can not account for all these missing wasps from his study, and he is inclined to believe that they out-migrated far enough away to preclude any further encounters at the site of their emergence (pers. comm. E-mail 29 March 2021),

I tend to believe that something similar occurred with the *S. pensylvanicus* at my study site; the female wasps may not necessarily be rigidly loyal to their natal location based on a few of my observations as discussed below.

First, looking at Table 5, one finds that in 2013 only six female *S. pensylvanicus* were caught from the apertures of the NLLRW. Then in 2014 and 2015, no wasps at all were caught emerging from the NLLRW apertures probably indicating the extinction of the aggregation. However, Table 5 also shows that in 2016 wasps again emerged from the apertures meaning that apparently some wandering female *S. pensylvanicus* had re-established a nesting aggregation in the wall during the previous season which I had not noticed. In 2017, even more *S. pensylvanicus* emerged from the apertures indicating further successful nesting during the 2016 season.

I also observed some possible exploring behavior in 2004 as evidenced by Wasp #57 from that year. She was initially captured by sweep net while flying in close proximity to Blocks 5 and 6 on 18 July. Then on 19 July Wasp #57 was noted making a visit of several seconds duration into a drain hole in the driveway retaining wall of the residence directly across the street from my lot. My last encounter with Wasp #57 was on 20 July; she was exiting from the 1E aperture of the NLLRW at that time.

Also, in 2004, another instance of an apparent wandering female *S. pensylvanicus* can be cited. On 21 July a female was captured from the 1W trap and given tag #62. On 1 August I saw a previously tagged female making long orientation flights over the front lawn of the residence at 3620 Virginia and directed at my <u>south</u> lot line retaining wall. This wasp was netted and found to be Wasp #62 and then released. Later, on 4 August I found Wasp #62 captured in the trap jar in the 1W aperture. This is noteworthy since the previously observed orientation flights on 1 August 2004 would seem to indicate that Wasp #62 had a nest in the south lot line retaining wall as well as possibly in the NLLRW (through the 1W aperture). While not absolutely proven in this case, could she have been tending to two different nests around the same time? I have found no records in the literature of *S. pensylvanicus* tending to more than one nesting location, although Baerends (1947) reported multiple nest tending by the ammophiline sphecid *Ammophila campestris* in Europe.

On a final note, Dr. Kurczewski has informed me that he can not confirm absolutely that the aggregation of *Sphex pensylvanicus* nesting in a storm drain in New York (Kurczewski 1997, 1998) were all emergers from this natal location; some of them could have flown in from elsewhere (pers. comm. E-mail 10 June 2017).

### Protandry and Territoriality

As previously stated, the emergence of the majority of males several days before the females (protandry) for *Sphex pensylvanicus* appears to be confirmed for the aggregation at this study site (See Table 5). If the purpose of protandry is to ensure that the males are afforded the best chance to fertilize the later emerging females, then a puzzling situation has presented itself at this location. From 2004 through 2017, I can verify 131 males as having emerged from the apertures of the NLLRW yet not one of these males established a territory on this wall, and I have no clue as to where they went nor what happened to them. Furthermore, it was only in 2003, 2004, and 2005 that I saw any males behaving in a territorial manner – a handful of males in 2003 and only one male each in both 2004 and 2005 (See Territoriality). Although incestuous mating among the Hymenoptera is frequent (See O’Neill, esp. Chapter 8), perhaps with *Sphex pensylvanicus* the disappearence of the males is an adaptation to reduce inbreeding.

## Acknowledgments

Many thanks to Dr. F.E. Kurczewski, State University of New York, Syracuse, New York, and Dr. H.J. Brockmann, University of Florida, Gainesville, Florida, for information and comments they provided on very early portions of this study. I am also indebted to Dr. W.J. Pulawski, California Academy of Sciences, San Francisco, California, for identification of the wasp *Liris argentatus*. I am also grateful to Dr. Carlo Polidori, Dipartimento di Scienze e Politiche Ambientali – ESP, Universita degli Studi di Milano, Milan, Italy, for providing information on an aggregation of the digger wasp *Stizus continuus* that he studied in Spain.

## Literature Cited

1. Baerends, G.P. 1947. On the Life-History of Ammophila campestris Jur. Proceedings of the Section of Sciences. Koninklijke Akademia van Wetenschappente Amsterdam, Vol. 44. pp. 483–488.

2. Bohart, R.M. and A.S. Menke. 1963. A reclassification of the sphecinae with a Revision of the Nearctic species of the tribes Sceliphronini and Sphecini (Hymenoptera: Sphecidae). University of California Press. Berkeley. 181 pp.

3. Buck, M. 2004. An annotated checklist of the spheciform wasps of Ontario (Hymenoptera: Ampulicidae and Crabronidae). Journal of the Entomological Society of Ontario. 134: 19–83.

4. Davis, W.T. 1920. Mating habits of Sphecius speciosus, the cicada killing wasp. Bull. Brooklyn Entomol. Soc. 15: 128–129.

5. Evans, H.E. 1966(a). The Behavior Patterns of Solitary Wasps. Annual Review of Entomology. 11: 123–154.

6. Evans, H.E. 1966(b). The Comparative Ethology and Evolution of the Sand Wasps. Harvard University Press. Cambridge. Massachusetts. 526 pp.

7. Frisch, J.A. 1938. The Life-history and Habits of the Digger-wasp, Ammobia pennsylvanica (Linn.) The American Midland Naturalist. 19: 673–677.

8. Kurczewski, F.E. 1997. Activity Patterns in a Nesting Aggregation of Sphex pensylvanicus L. (Hymenoptera: Sphecidae). Journal of Hymenoptera Research. Vol. 6(2). pp. 231–242.

9. Kurczewski, F.E. 1998. Territoriality and Mating Behavior of Sphex pensylvanicus L. (Hymenoptera: Sphecidae). Journal of Hymenoptera Research. Vol. 7(1). pp. 74–83.

10. Lechner, G.K. 2015. Large Homing Flight Distances Flown by Female Sphex pensylvanicus, Linnaeus, 1763 (Hymenoptera: Sphecidae: Sphecinae) in Sioux City, Iowa, U.S.A. Life: The Excitement of Biology. 3(2). Pp.149–152.

11. Lechner, G.K. 2016. Interesting Incidents with Sphex pensylvanicus Linnaeus, 1763 (Hymenoptera: Sphecidae) Wasps and Their Prey Items in Sioux City, Iowa, U.S.A. Life: The Excitement of Biology. 4(1). pp. 27–31.

12. Lewis, J.H. 2020, Sphex ichneumoneus and Sphex pensylvanicus (Hymenoptera: Sphecidae) in Atlantic Canada: evidence of recent range expansion into the region. The Canadian Field-Naturalist. Vol. 134. pp. 52–55.

13. Lin, N. 1963. Territorial Behavior in the cicada killer wasp Sphecius speciosus (Drury) (Hymenoptera: Sphecidae). Behavior 20: 155–133.

14. O’Neill, K.M. 2001. Solitary Wasps Behavior and Natural History. Ithaca and London. Cornell University Press. 406 pp.

15. Polidori, Carlo, Irene Giordani, Pablo Mendiola Josep, D. Asís, Jose Tormes, Jesús Selfa. 2010. Emergence and dispersal relative to natal nest in the digger wasp Stizus continuus (Hymenoptera: Crabronidae). Comptes Rendus Biologies. 333: 255–264.

16. Rau, P. 1944. The Nesting Habits of the Wasp, Chlorion (Ammobia) pennsylvanica L. Annals of the Entomological Society of America. 37: 439–440.

17. Rigley, L. and H. Hays. 1977. Field Observations Including Acoustic Behavior of the Black-Digger Wasp, Sphex pennsyvanicus (Linn.). Proceedings of the Pennsylvania Academy of Science. 51: 32–34.

